# Dietary fibre promotes chronic whipworm infection through direct and time-dependent modulation of innate immunity

**DOI:** 10.1101/2025.06.06.658267

**Authors:** Laura J. Myhill, Penille Jensen, Pankaj Arora, Anne M. Jensen, Ling Zhu, Amalie Vedsted-Jakobsen, Eiríkur A. Thormar, Alexandra von Munchow, Mahesha M. Poojary, Marianne N. Lund, Stig M. Thamsborg, Morten T. Limborg, Benjamin A. H. Jensen, Andrew R. Williams

## Abstract

Dietary fibre regulates the microbiome and gut health but increases murine whipworm (*Trichuris muris*) infection through unclear mechanisms. We show that mice fed inulin-supplemented diets exhibit dysregulated innate antimicrobial defences and altered tryptophan metabolism during *T. muris* infection. Inhibiting tryptophan catabolism or neutralizing IL-27 and IL-18 in inulin-fed mice restored infection resistance. Notably, inulin led to chronic infection even in microbiota-depleted mice. Removing inulin within a critical immune development window rapidly restored anti-helminth immunity, indicating direct, time-dependent modulation of mucosal immune responses. Our findings reveal a previously unrecognized, direct influence of dietary fibre on mucosal immunity to parasitic infection, independent of the microbiome, highlighting the complex interplay between diet timing and host defence.

## Introduction

Diet composition profoundly influences mucosal immunity and host defence against pathogens ^1^. Dietary components, including macro- and micronutrients and dietary fibre, can directly modulate immune cell function or indirectly impact immunity by shaping the gut microbiota (GM) and its metabolic output^2, 3^ ^4, 5, 6^.

Modern diets, often characterized by a deficit in unprocessed plant material and fibre, promote gut dysbiosis, chronic inflammation and increased infection susceptibility^2, 7^. Consequently, fermentable fibres like inulin are increasingly used as dietary supplements to restore gut homeostasis^8, 9, 10, 11^. The mode-of-action of bioactive dietary substances such as inulin is highly complex. As a fermentable fibre, inulin intake invariably results in drastic changes in GM composition and increased production of bacterial-derived metabolites such as short chain fatty acids (SCFA), and effects of inulin on gut health and disease outcomes are often ascribed to these changes^12^. However, inulin (a large fructan-based polymer) also interacts directly with mammalian cells and can induce immunological changes in the gut of germ-free as well as conventional mice^13, 14^, suggesting that multiple overlapping mechanisms may contribute to the biological activity of inulin.

Paradoxically, inulin can also exacerbate certain diseases, including colitis and liver cancer in mice^15, 16^. Aggravation of irritable bowel syndrome can also occur in humans following inulin consumption^17^. Moreover, we have previously demonstrated that incorporation of inulin (but not the insoluble fibre cellulose) into purified mouse diets increases susceptibility to the parasitic whipworm *Trichuris muris* (*Tm*).*Tm* is the murine equivalent of the human pathogen *Trichuris trichiura* which infects over 400 million people and causes significant morbidity, including growth retardation, malnourishment, and chronic intestinal disease^18, 19^. Immunity to *Tm* relies on type-2 immune responses^20^, but inulin-fed mice fail to mount protective immunity, exhibiting type-1 biased inflammation, suggesting that the level of dietary fibre acts to regulate the balance between opposing immune cell subsets during intestinal parasite infection^18, 21^.

Given the prevailing view that the immunomodulatory effects of inulin are primarily mediated by the GM, we investigated the mechanisms underlying inulin-induced susceptibility to *Tm*, focusing on the interplay between the GM, host immunity, and parasite interaction. Unexpectedly, we discovered that inulin directly modulates innate immunity, independently of the GM, via a time-dependent mechanism involving IL-27 and IL-18. This work uncovers a novel, direct pathway by which dietary fibre drives susceptibility to parasitic infection, challenging the dogma of exclusive microbiome-mediated effects and offering new insights into the diet-host-pathogen axis.

## Results

### Dietary inulin and Trichuris muris infection modulate local cellular responses in the large intestine

We previously demonstrated that dietary inulin consumption during *Tm* infection resulted in increased worm burdens, intestinal inflammation, and enhancement of adaptive T helper (Th)-1 responses in mesenteric lymph nodes (MLN) at 21 days post-infection (p.i.)^18^. To explore whether these outcomes derived from a defect in the early immune response elicited by *Tm*, we first assessed innate and adaptive cellular responses in large intestinal lamina propria (LP) of mice fed either purified, semi-synthetic diets (SSD) or inulin-enriched SSD (hereafter ‘inulin’; Supplementary Table 1), with or without 7 days of *Tm* infection (**Figure 1a**). Dietary inulin and *Tm* infection independently increased both the number and proportion of CD4^+^ T-cells (*p*<0.001 for main effects of diet and infection; **Figure 1b**). Within the CD4+ T-helper population, *Tm* infection significantly increased the proportions of Th1 (Tbet^+^) and Th2 (GATA3^+^) cells, with Th1 cells being the dominant population at this time-point. However, these expansions were equivalent in both diets, suggesting that inulin did not alter the adaptive immune polarization early in the infection (*p*<0.01 for main effect of infection; **Figure 1c-d**). Interestingly, *Tm* infection significantly decreased the proportion of Th17 cells (**Figure 1e**), whilst simultaneously boosting the Tbet^+^ subset within this cell population (Supplementary Figure 1). However, these changes were, akin to the Th1 and Th2 profile, only affected by *Tm* infection and not diet. Moreover, innate lymphoid cell (ILC) sub-populations (ILC1, ILC2 and ILC3) in the large intestine LP, and dendritic cell populations in mesenteric lymph nodes, were not significantly altered by either infection or diet (Supplementary Figure 1). Taken together, these data indicate that *Tm* elicited a mixed Th1 and Th2 cellular response, concomitant with a decreased and aberrant Th17 response, in the intestinal mucosa early in infection. Furthermore, despite a drastic inulin-dependent Th1 bias later in the infection^18^, inulin only affected the numbers of CD4^+^ T cells but not their polarization in the early T-cell response.

**Figure 1.**
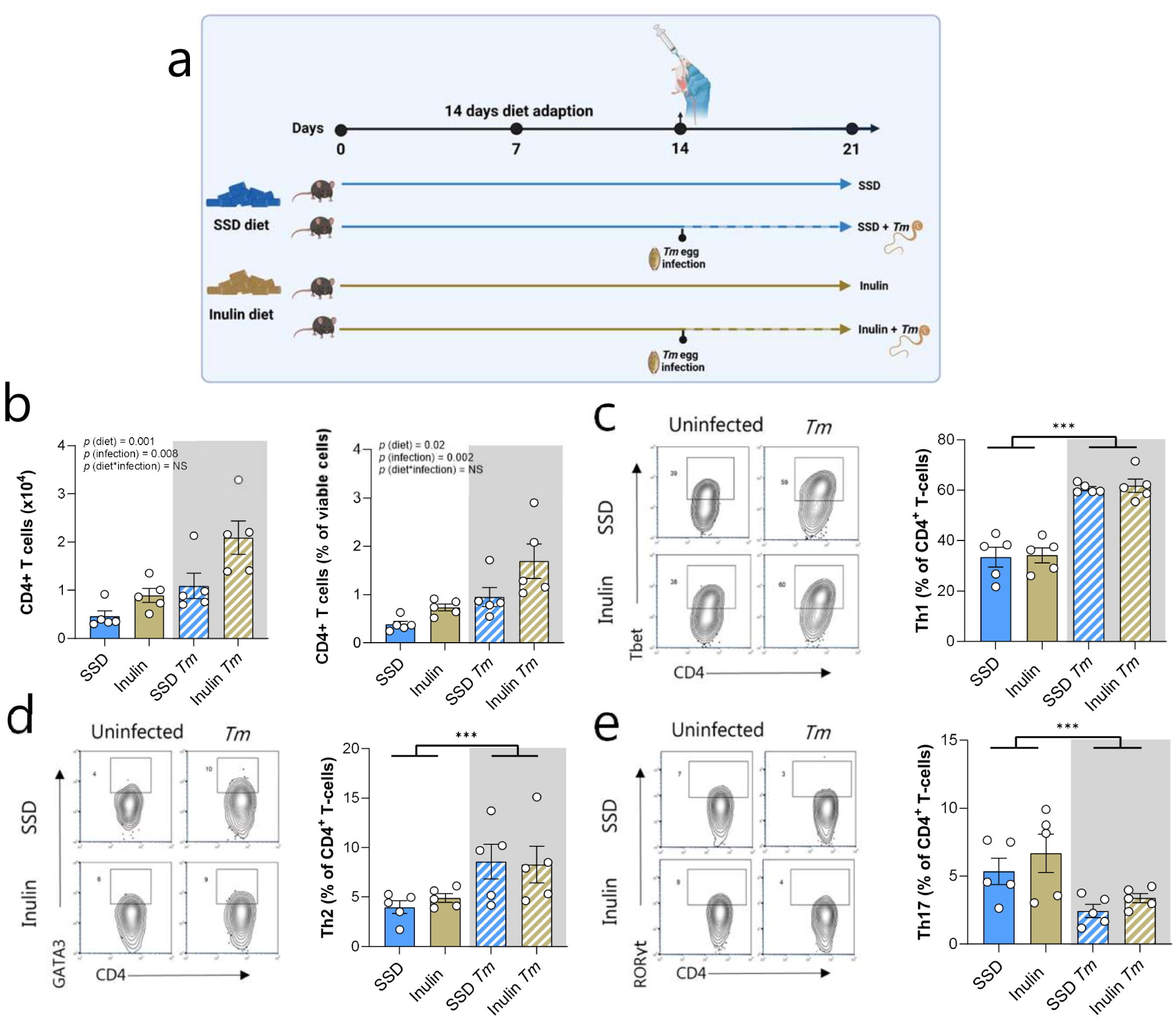
Dietary inulin and *Trichuris muris* infection modulate T-cell populations in the large intestine. **a)** Experimental design – mice were fed either SSD or SSD supplemented with 10% inulin (see methods). After two weeks of diet adaptation, half the mice in each dietary group were infected with 300 *Trichuris muris* (*Tm*) eggs for 7 days. **b)** Numbers and proportions of CD4^+^ T-cells in large intestinal tissue. Proportions of Th1 (**c**), Th2 (**d**) and Th17 (**e**) in CD4^+^ T-cells in large intestinal tissue. n=5 per treatment group. Data shown are means ± S.E.M. *** *p*<0.001 indicates main effect of infection by two-way ANOVA. See Supplementary Figure 6 for gating strategy.

### Dietary inulin dysregulates innate epithelial responses to Trichuris infection

The intestinal epithelium is the first site of contact between *Tm* and the host. Release of alarmins (especially IL-25, IL-33, and TSLP) from intestinal epithelial cells (IEC) in response to larval invasion is critical for subsequent development of protective immune responses^22^. Whilst we observed no differences between diets in the ILC or early T-cell response, we speculated that inulin-driven alterations in IEC function may drive inflammation that progressively supresses the development of the protective immune response and worm expulsion. At day 3 p.i, *Il33, Tslp,* and *Ifng* expression in whole caecal tissue was not changed by diet or *Tm* infection (data not shown), but at day 7 p.i., expression of all three genes was markedly increased in infected mice, independently of diet (**Figure 2a**).

**Figure 2.**
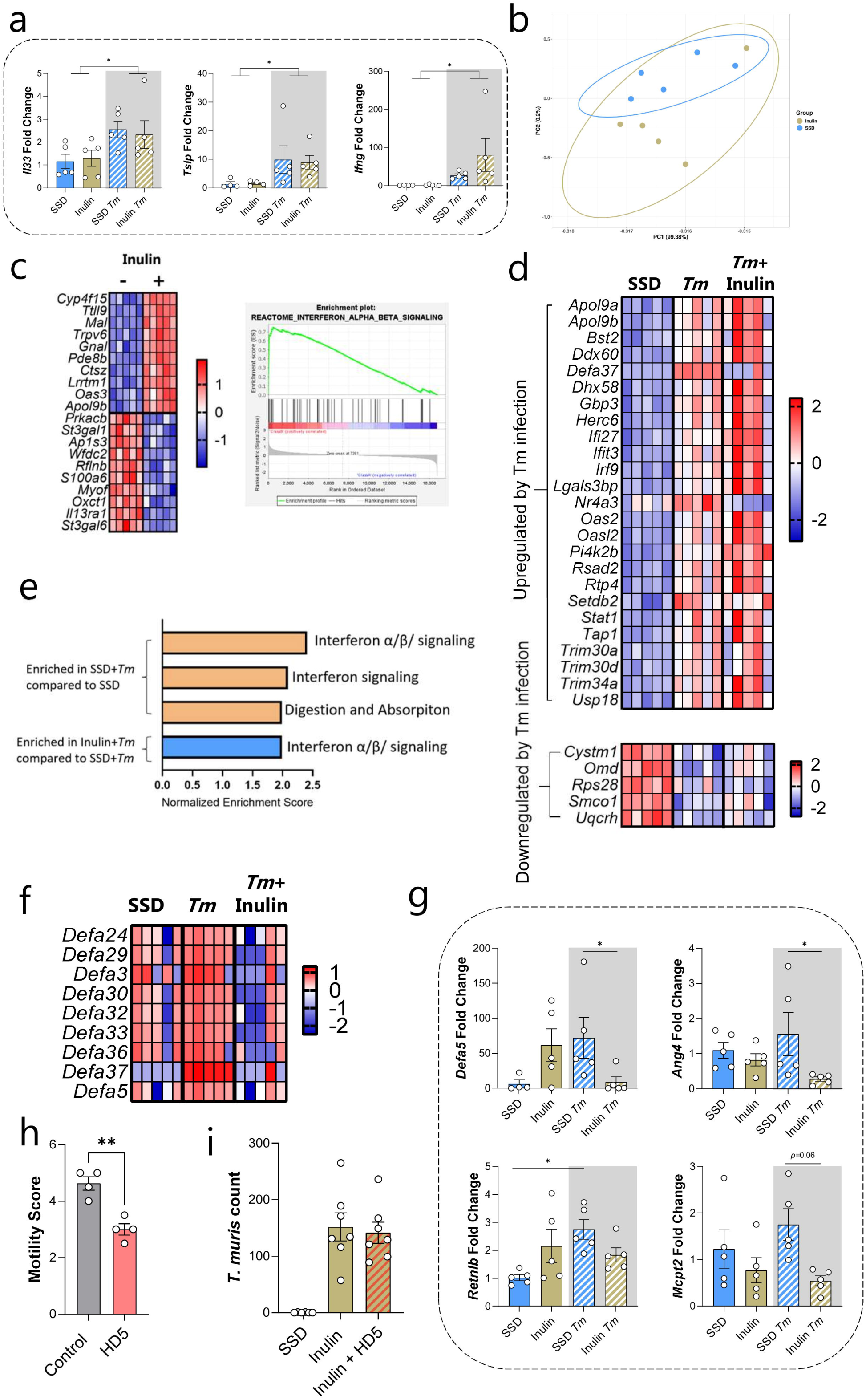
Dietary inulin influences the innate response to *Trichuris muris*. **a)** *Il33, Tslp* and *Ifng* expression in caecal tissue 7 days after infection with 300 *Trichuris muris* (*Tm*) eggs in mice fed either SSD or inulin. **b)** Principal component analysis of the intestinal epithelial cell (IEC) transcriptome in uninfected mice fed either SSD or inulin for 21 days. **c)** Top ten up- and down-regulated genes (adjusted *p*<0.05 by DESeq2) identified by RNA-Seq (shown are Z-scores), and enrichment of REACTOME interferon alpha/beta pathway, in IEC of uninfected mice fed either SSD or inulin for 17 days. **d)** Genes identified by RNA-Seq (Z-scores) as being significant regulated (adjusted *p* value <0.05 using DESeq2) by *Tm* in IEC of SSD-fed mice 3 days post-infection, and their corresponding expression in inulin-fed mice infected with *Tm*. **e)** Pathways identified by gene-set enrichment analysis as being significantly (*q*<0.05) enriched by *Tm* infection in SSD-fed mice, and by inulin diet relative to SSD diet within *Tm*-infected mice. **f)** Expression (Z-scores) of genes encoding α-defensins in mice fed SSD, SSD with *Tm* infection, or inulin with *Tm* infection. **g)** qPCR analysis of *Ang4, Defa5, Retnlb* and *Mcpt1* expression in IEC in mice fed SSD or inulin, with or without 3 days of *Tm* infection. **h)** Motility of *Tm* larvae exposed for 24 hours to recombinant human defensin-5 (HD-5). Motility was scoured by a blinded observer as indicated in methods. **i)** *Tm* burdens 21 days post-infection, in mice fed either SSD, inulin, or inulin together with daily HD-5 treatment between day 0 and day 21 p.i. Data shown are means ± S.E.M. **a) – g)** n = 5 per group, **i)** = 6 per group. **a)** * *p*<0.05 for main effect of infection by two-way ANOVA. **g)** * *p*<0.05 by two way ANOVA and Tukey post-hoc testing. **h)** ** by two-tailed t-test.

We next isolated IEC cells from the large intestine (combined caecum and colon) at day 3 p.i, in mice fed inulin or SSD, and their uninfected controls. RNAseq analysis of IEC showed that in the absence of infection, inulin markedly altered the epithelial transcriptome (**Figure 2b**). Downregulated genes included *Oxct1*, encoding a transferase involved in ketolysis, and the IL-13 receptor alpha 1 subunit (*Il13ra1*). Notably, inulin resulted in the significant upregulation of numerous genes encoding molecules involved in antimicrobial responses such as, interferon-induced proteins (*Ifit* gene family) and 2’-5’-linked oligoadenylate synthase enzymes (*Oas* gene family) (**Figure 2c**; Supplementary File 1), which was also reflected in significant enrichment of the REACTOME interferon signalling pathway by gene set enrichment analysis (**Figure 2c**). Thus, inulin (in the absence of any pathogen infection) appeared to induce a heightened state of IEC-mediated immune reactivity in the large intestine. In SSD-fed mice, *Tm* enhanced expression of numerous genes involved in innate immune defences, e.g. interferon-induced proteins (*Ifit3*, *Ifi27*, *Irf9*), and *Oas* family genes, as well as genes involved in metabolic processes such as lipid and amino acid metabolism (**Figure 2d**). Pathway analysis revealed interferon signalling and antiviral immune pathways to dominate these responses, corroborating recent *in vitro* results from mouse caecaloids^23^ **(Figure 2e**). The expression of genes involved in interferon signalling, e.g. *Usp18* as well as the *Ifit* and *Oas* gene families, tended to be enhanced in infected mice fed inulin (**Figure 2d**). Consistent with this, gene pathway analysis revealed that the interferon signalling pathway was significantly enhanced in infected mice fed inulin, relative to infected mice fed SSD (**Figure 2e**), indicating an additive effect of inulin and *Tm* in inducing an interferon rich signalling environment. Interestingly, we noted significant upregulation of *Defa37* and *Defa5*, encoding members of the α-defensin family, in *Tm*-infected mice. α-defensins are produced mainly from Paneth cells which are present in the small intestinal crypts. However, Paneth cell metaplasia in the colon and associated α-defensin production is a feature of large intestinal pathogens, including *Tm*^24^. Notably, the expression of both *Defa37* and *Defa5* was markedly attenuated in *Tm*-infected mice upon inulin feeding (**Figure 2d**), suggesting an aberrant α-defensin response to infection. *Defa5* (and *Defa24* which was also nominally decreased in inulin fed mice) has previously been implicated in immunity against *Tm*^25^ (**Figure 2f**).

We confirmed by qPCR that the expression of *Defa5*, despite being upregulated by both inulin and *Tm* infection alone, was completely abrogated in infected mice fed the inulin diet (**Figure 2g**). To explore if other innate effector peptides were also modulated by dietary inulin, we measured the expression of *Ang4, Retnlb* and *Mcpt2* which showed a similar pattern, i.e. the expression of these was induced by infection in SSD-fed mice, but attenuated in infected, inulin-fed mice, suggesting a profound defect in the innate immune response to *Tm* (**Figure 2g**). To explore the functional implications of the impaired α-defensin production, we treated inulin-fed, *Tm-*infected mice with recombinant human α-defensin-5 (HD-5) to interrogate if this would restore innate antimicrobial activity and aid resistance to *Tm.* As has been previously shown with recombinant murine Defa5^25^, *Tm* worms cultured *ex vivo* with HD-5 displayed a noticeable reduction in motility (**Figure 2h**). Nevertheless, inulin-fed infected mice treated with HD-5 had similar worm burdens compared to inulin fed mice without HD-5 treatment (**Figure 2i**), indicating that the magnitude of innate immune impairment surpassed the rescue-potential of a stand-alone treatment. Collectively, these data indicate that as early as 72 hours after infection, dietary inulin results in a profound remodelling of the epithelium transcriptional response to infection with impaired production of antimicrobial peptides, but that restoration of a select α-defensin response was not sufficient to re-establish immunity and worm expulsion.

### Modulation of host tryptophan or AhR signalling partially restores immunity to Trichuris muris in inulin-fed mice

We next asked how these dysregulated innate responses early in *Tm* infection led to suppression of the protective anti-helminth response in inulin-fed mice. We previously demonstrated that 21 days after *Tm* infection the caecal transcriptome in inulin-fed mice became dominated by expression of genes involved in type-1 inflammation such as *Ifng* and *Cxcl10*, but that the most over-expressed gene was *Ido1,* encoding the enzyme indoleamine 2,3-dioxygenase 1 (IDO1), which is responsible for the catabolism of dietary tryptophan to kynurenine^18^. IDO1 expression is induced by an interferon-rich environment, and, importantly, has been previously shown to mitigate expulsion of *Tm*^26^. Given that the interferon signalling pathway was the only transcriptional pathway to be enriched in inulin-fed mice 3 days p.i., relative to SSD-fed mice, we postulated a role for IDO1-mediated altered tryptophan metabolism in mediating *Tm* infection. To explore this, we first assessed serum tryptophan and kynurenine concentrations during *Tm* infection, which revealed significantly increased kynurenine:tryptophan ratios in inulin-fed mice compared to SSD-fed mice, suggesting a higher rate of tryptophan catabolism (**Fig. 3a**). Moreover, histochemical staining of proximal colon tissue revealed IDO1 to be substantially higher in inulin-fed mice compared to SSD-fed controls (**Fig. 3b**). To assess the functional importance of IDO1 signalling, infected mice fed the different diets were treated with the IDO1 inhibitor 1-methyl-DL-tryptophan (1-MT) or vehicle control. Notably, 1-MT treatment partially restored *Tm*-induced goblet cell hyperplasia that was severely inhibited in inulin-fed mice (*p*<0.05 for main effects of diet and 1-MT treatment; **Fig. 3c**). Worm counts were also lower in 1-MT treated mice, regardless of diet, showing that immunity was partially restored (**Fig 3d**). qPCR analyses in caecal tissue showed that 1-MT also partially abrogated and boosted the inulin-induced increase in *Ifng* expression and *Il13* expression, respectively (**Fig 3e**).

**Figure 3.**
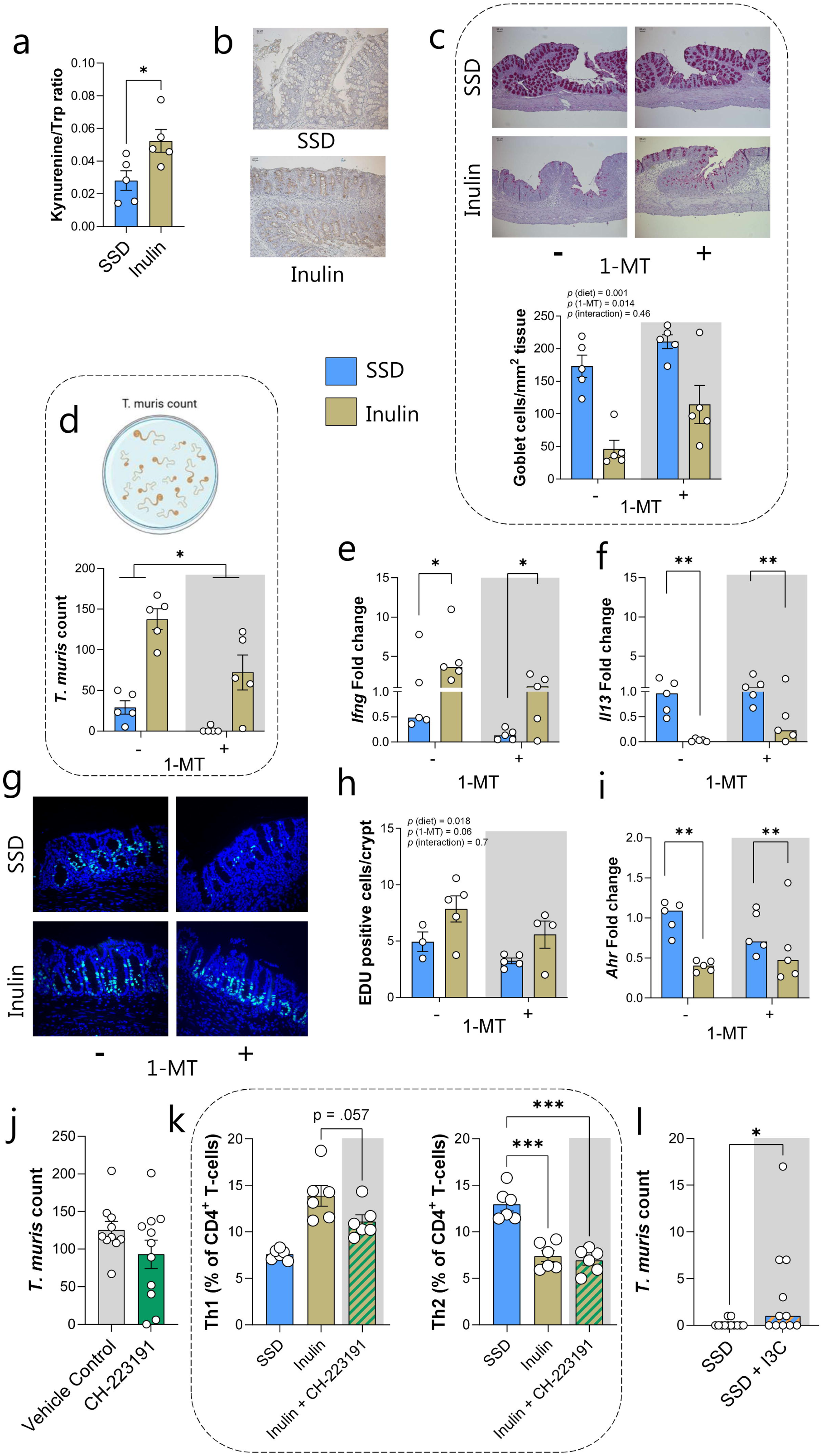
Pharmacological manipulation of tryptophan metabolism or Aryl Hydrocarbon Receptor signaling regulates *Trichuris muris* burdens in inulin-fed mice. **a)** Ratio of kynurenine to tryptophan (Trp) in serum of *Trichuris muris* (*Tm*)-infected mice at day 21 p.i., fed either SSD or inulin diets. **b)** Representative images of the proximal colon of *Tm-*infected mice at day 21 p.i., fed either SSD or inulin diet, showing immunohistochemical detection of IDO1. **c)** Representative images of PAS staining and goblet cell counts in proximal colon, **d)** *Tm* burdens, **e)** *Ifng* and *Il13* expression in caecum, **f)** Representative images of EDU staining and positive cell counts in proximal colon, and **g)** *Ahr* expression in caecum at day 21 of *Tm* infection in mice fed either SSD or inulin diet and administered either 1-MT or vehicle control in drinking water. **h)** *Tm* burdens (day 21 p.i.) in mice fed inulin diet and administered daily treatment with vehicle control or AhR antagonist CH-223191. **i)** proportions of Th1 (Tbet^+^) and Th2 (GATA3^+^) CD4^+^ T-cells in the mesenteric lymph nodes of mice infected with *Tm* (day 21 p.i.) and fed either SSD, inulin + vehicle control, or inulin + CH-223191. **j)** *Tm* burdens (day 21 p.i.) in mice fed either SSD or SSD + 200mg/kg indole-3-carbinol (I3C). Data shown are means ± S.E.M except **e)** and **g)** for which medians are shown. **a)** and **j)** *p<0.05 by two-tailed t-test. **c)** and **f)** *p<0.05;**<0.01, ***<0.001 by two-way ANOVA and Tukey post-hoc testing. **d)** * *p*<0.05 for main effect of 1-MT treatment by two-way ANOVA. **e)** **p<0.01 for main effect of diet by two-way ANOVA. **i)** ***p<0.001 by one-way ANOVA and Tukey post-hoc testing. **a-g and i) –** n=5 from one experiment. **h and j) –** pooled from two independent experiments with 5-6 mice per group/experiment.

IEC homeostasis is under tight control of both cytokines and nutrient- and metabolite-sensing pathways, including Trp catabolism^27^. We noted that colonic crypt lengths were higher in inulin-fed mice during *Tm* infection, and that there were more Edu^+^ positive cells in the crypts indicative of increased proliferation - a hallmark of *Tm* chronicity^28^ – with these effects being somewhat muted by 1-MT treatment (*p*<0.05 for main effect of diet, *p*=0.06 for main effect of 1-MT; **Figure 3f**). Breakdown of Trp by the host via IDO1 to kynurenine, or by gut bacteria to yield indole derivatives such as indole-3-lactic acid, activates the Aryl Hydrocarbon Receptor (AhR) to regulate IEC activity^29^. Interestingly, we found that expression of *Ahr* expression was significantly suppressed in inulin-fed mice (but not modulated by 1-MT treatment) (**Figure 3g**). This may suggest that the AhR pathway was perturbed by dietary inulin. Interestingly, AhR signalling has been shown to regulate anti-helminth immunity, with AhR-deficient mice being more resistant to infection with the roundworm *Heligmosomoides polygyrus*^30^, which prompted us to test a role for the AhR pathway in mediating susceptibility to *Tm.* Inulin-fed mice were administered daily the AhR antagonist CH-223191, or vehicle control, during *Tm* infection. Infection was not significantly reduced, although we did note that 2 mice given the antagonist had cleared or nearly cleared their infection (0 and 6 worms, respectively; **Figure 3h**), something not observed in any of our other experiments with inulin-fed mice, where worm burdens were typically in the range of 100-150 worms. Moreover, CH-223191-treated mice tended to have lower proportions of Th1 cells in their mesenteric lymph nodes, with Th2 proportions unchanged (**Figure 3i**). We then tested whether mice fed SSD (essentially devoid of dietary AhR ligands) would have increased *Tm* burdens when the SSD was supplemented with indole-3-carbinol (I3C), a prodrug which is metabolized in the stomach to highly potent AhR agonists such as 5,11-dihydroindolo-[3,2-b]carbazole. Mice fed SSD with I3C had significantly higher *Tm* burdens than their non-I3C fed counterparts, however still with very low numbers (range 0 to 17 worm; **Figure 3j**). Overall, these experiments show that modulation of either the host Trp or AhR signalling pathways could alter the inulin-induced immune responses and *Tm* burdens but were not sufficient to recapitulate the marked effect of dietary inulin.

### IL-27 and IL-18, but not IL-6, drive pro-inflammatory responses and susceptibility to Trichuris muris in inulin-fed mice

As 1-MT treatment was only partially effective in restoring immunity to *Tm* in inulin-fed mice, we postulated that interventions acting upstream of IDO1 activity would be more effective. In addition to release of alarmins from IEC, innate cytokines released from macrophages or DCs are critical factors in determining the initial balance of immune molecules that instruct the type-2 response and anti-helminth immunity^31^. Specifically, IL-27 (also known as WSX-1), IL-6, and IL-18 have been implicated in inhibiting immunity to *Tm*^32, 33, 34, 35^. Inulin-fed mice infected with *Tm* were given neutralizing anti-IL-6, anti-IL-27, or anti-IL-18 antibodies during the early stages of infection. Strikingly, blocking IL-27 resulted in almost complete restoration of *Tm* expulsion (**Figure 4a**), and IL-18 blockage also substantially reduced worm burdens (**Figure 4b**), with anti-IL-6 treatment having no effect (**Figure 4c**). Intriguingly, IL-27 and IL-18 are also interferon-responsive cytokines having a key role in IDO1 upregulation in inflammatory settings^36, 37^. Anti-IL-27 treatment efficiently supressed the expression of both *Ifng* and *Ido1* in caecum tissue at day 21 p.i. (**Figure 4d-e**), and anti-IL-18 treatment also tended to reduce *Ido1* expression without affecting *Ifng* (**Figure 4f-g**). These data confirm that an altered innate response early in infection, driven primarily by IL-27 and IL-18, promotes *Tm* infection and that this may involve Th1 polarization and altered *Trp* metabolism.

**Figure 4.**
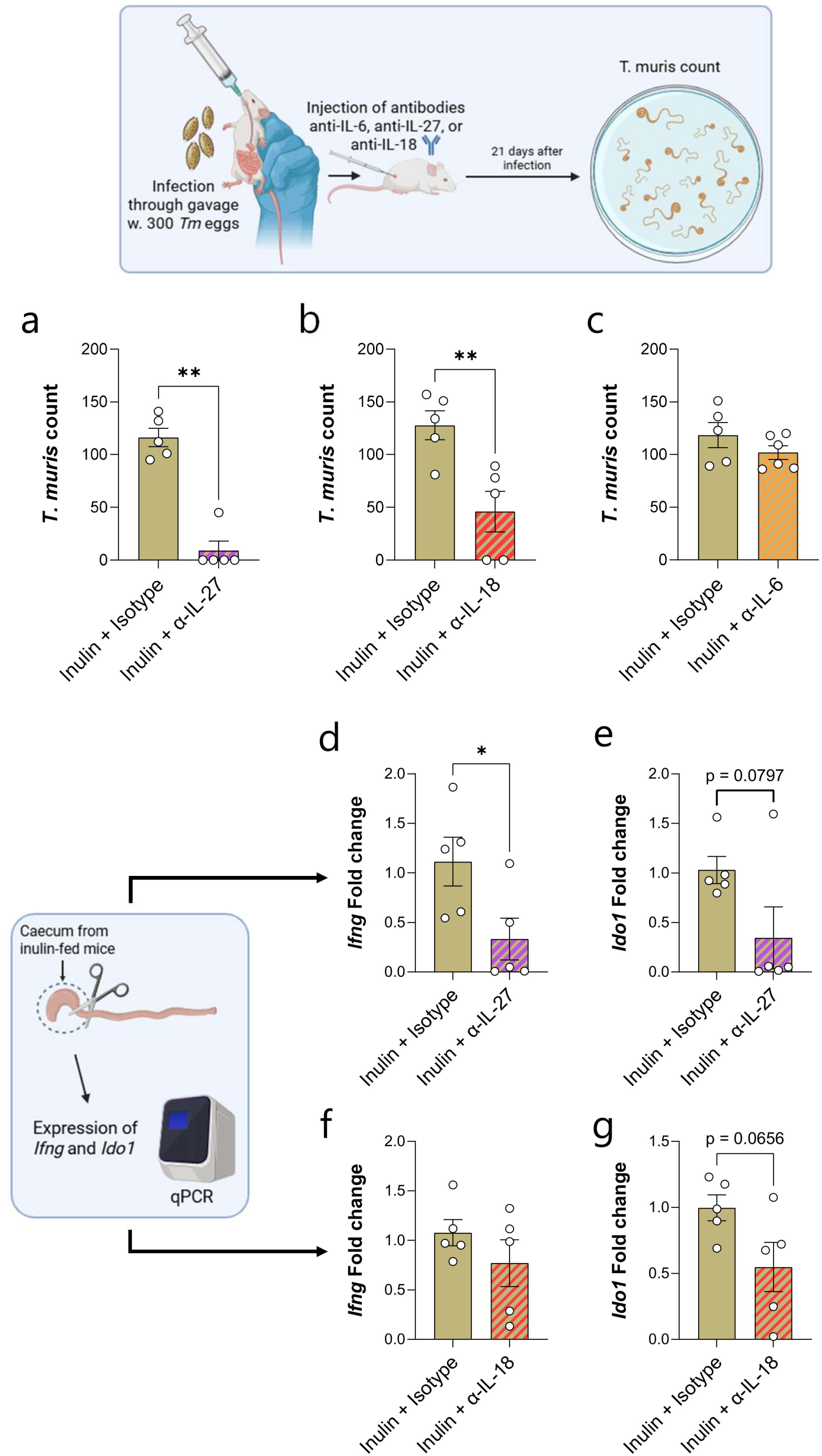
Neutralization of selected innate cytokines restores immunity to *Trichuris muris* in inulin-fed mice. **a-c)** *Trichuris muris* burdens in mice fed inulin diet, 21 days after infection with 300 eggs, and administered either isotype controls or neutralizing antibodies against IL-27 (**a**), IL-18 (**b**) or IL-6 (**c**). Data shown are means ± S.E.M (IL-6, IL-18) or medians (IL-27). ** *p*<0.01 by Mann-Whitney test (IL-27) or unpaired t-test (IL-18). **d-g)** Expression of *Ifng* and *Ido1* in caecum tissue in inulin-fed mice infected with *Tm*, and administered either isotype controls or neutralizing antibodies against IL-27 (**d-e**) or IL-18 (**f-g**). Data shown are means ± S.E.M * *p*<0.05 by unpaired t-test. n=5 from a single experiment.

### *Trichuris muris* infection susceptibility is microbiota independent

As a highly fermentable fibre, breakdown of inulin by the GM promotes selective proliferation of many bacteria taxa and associated metabolite production. Indeed, we have previously shown that the mice consuming inulin have a drastically different GM composition than those fed SSD^18, 21^. To explore if this altered GM was a causative factor in the impaired immunity to *Tm*, we first assessed the role of SCFA, the main fermentation product of dietary inulin^38^. Inulin-fed mice were infected with *Tm* and treated with either an inhibitor of SCFA fermentation (β-acids from hops)^16^, or control drinking water. *Tm* burdens were not affected by inhibition of fermentation (**Figure 5a**). Moreover, addition of butyrate to the drinking water of either SSD- or inulin-fed mice also had no effect on worm burdens (**Figure 5a)**, suggesting that inulin-induced SCFA production is unlikely to play an important role.

**Figure 5.**
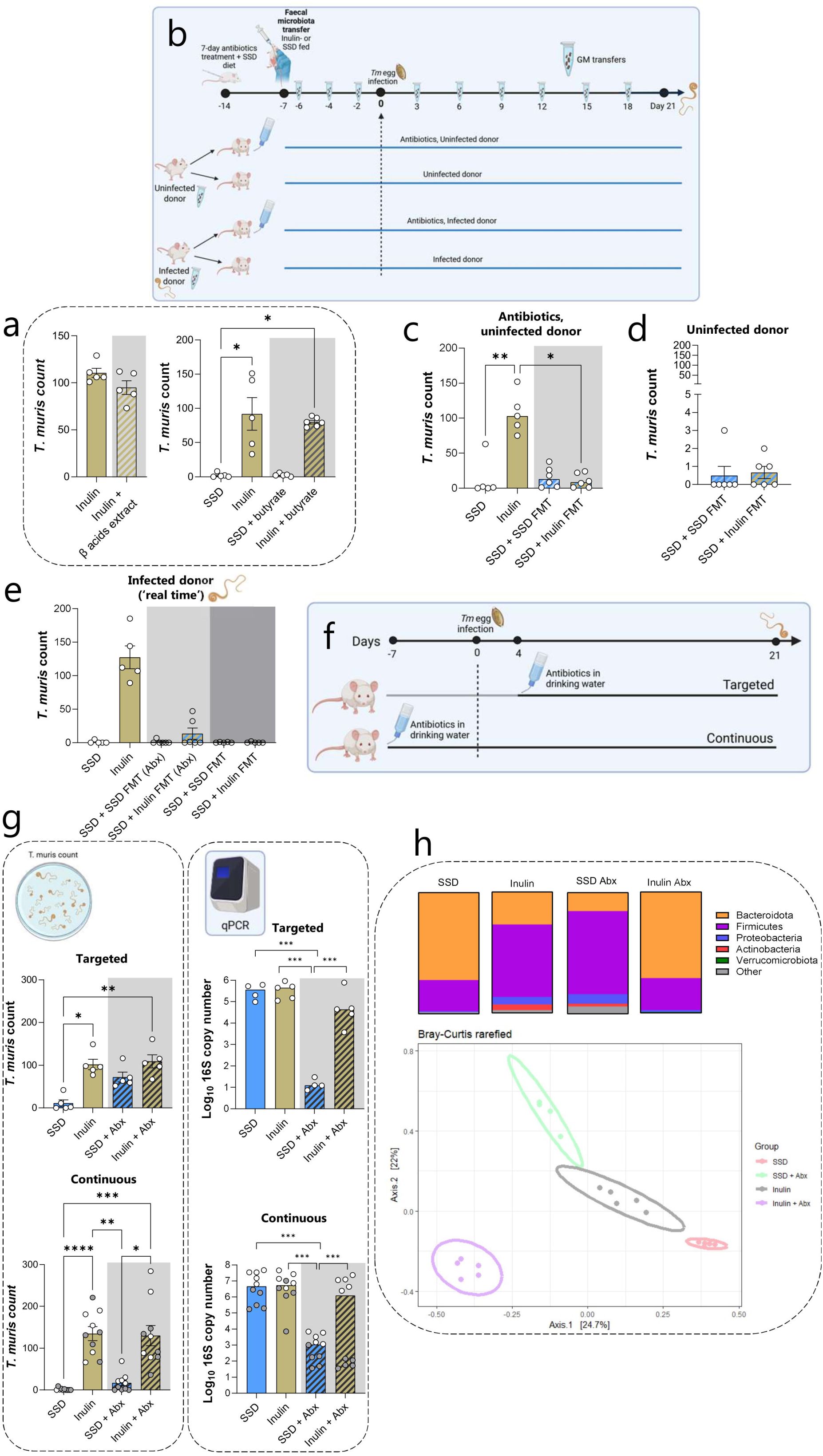
Effects of faecal microbiota transfer or antibiotic treatment on *Trichirs muris* burdens in inulin-fed mice. **a)** *Trichuris muris* (*Tm*) burdens at day 21 post-infection (p.i.) in SSD- or inulin-fed mice administrated butyrate or β-acids from hops. **b)** Design of faecal microbiota (FMT) studies. **c-e)** *Tm* burdens 21 days p.i. in SSD- or inulin-fed mice, or SSD-fed mice receiving FMT. **f)** Design of antibiotic administration studies. **g)** *Tm* burdens at day 21 p.i. and faecal 16S rRNA copy number in mice fed SSD or inulin, with or without targeted or continuous antibiotic treatment. For targeted antibiotic treatment, data are from a single experiment (n =5 per group). For continuous antibiotic treatment, data are from two independent experiments, each with n=5 per group, with the different experiments denoted by either open or closed circles. **h)** Faecal microbiota profile of *Tm*-infected mice fed SSD or inulin, with or without targeted antibiotic treatment. Data shown are distribution of bacterial phyla, and β-diversity analysis (PCoA plot of Bray-Curtis distance metrics).*p<0.05, **p<0.01, ***p<0.001 by one-way ANOVA and Tukey post-hoc testing.

Next, to scrutinize the role of inulin-induced GM populations, we employed a faecal microbiota transfer (FMT) approach. Mice were fed SSD and then depleted of their GM by a 7-day treatment with antibiotics (‘Antibiotics, uninfected donor’; **Figure 5b**). Mice where then recolonized with a faecal slurry (3 gavages) from uninfected mice fed either SSD or inulin, 7 days prior *Tm* infection. The GM transfers continued every three days after *Tm* infection until day 21 p.i. (**Figure 5b**). Mice receiving the inulin FMT were just as resistant as their SSD-colonized counterparts, suggesting a key influence of the inulin diet *per se* and not GM communities (**Figure 5c**). 16S rRNA amplicon sequencing of faecal samples from uninfected donor and recipient mice (prior to infection) indicated that both groups of mice receiving FMT formed a distinct cluster (likely due to prior antibiotic treatment). However, β-diversity analysis showed that the GM composition was different between these two FMT recipient groups (adjusted p-value <0.05 by PERMAMOVA on Bray-Curtis distance metrics), with their GM aligning more closely with their respective donor mice (Supplementary Figure 2). Notably, some of the major taxonomical difference between inulin- and SSD-fed donors (such as reduced *Akkermansia* spp. and increased *Bifidobacteria* spp. and *Bacteroides* spp. in inulin-fed mice) were recapitulated in mice receiving FMT from inulin-fed donors (Supplementary Figure 2). However, the GM of FMT-inulin recipient mice was, expectedly, significantly different to that of their donors (adjusted p-value <0.05 by PERMAMOVA on Bray-Curtis distance metrics), suggesting only partial engraftment of the donor GM in the absence of the prebiotic inulin substrate in the diet (Supplementary Figure 2).

To rule out a confounding effect of antibiotic treatment, we repeated the FMT studies in mice that received no prior antibiotic treatment (‘Uninfected donor’; **Figure 5b**). Again, SSD-fed mice that received FMT from either SSD-fed or inulin-fed mice were highly resistant to *Tm*, with infection levels of less than 5 worms (**Figure 5d**). Finally, to explore if progressive dysbiosis in inulin-fed mice during *Tm* infection leads to the impaired expulsion, we employed a dynamic ‘real-time FMT’ approach where inulin-fed mice were infected with *Tm* and a faecal slurry from these mice transferred every 3 days into SSD-fed mice at an equivalent time-point of infection (‘Infected donor’; **Figure 5b**). Again, these recipient mice were fully resistant to *Tm* infection both with and without antibiotic treatment prior to FMT (**Figure 5e**).

To further examine a role for the GM in mediating the dietary effect, mice fed SSD or inulin were infected with *Tm* and given a suppressive antibiotic cocktail (reported to deplete >90% of the mouse GM^16^) in drinking water. We employed two different experimental approaches. As antibiotic treatment may influence *Tm* hatching^39^, we first commenced antibiotic treatment 4 days after infection, to allow larvae to establish. In a second study, we administered antibiotics throughout the full experiment, from day 7 p.i. to day 21 p.i. (**Figure 5f**), hence mirroring the antibiotic regiment from our FMT studies (**Figure 5b-e**). Regardless of experimental approach, *Tm* burdens were equally high in inulin-fed mice (**Figure 5g**) and hence unlikely to be affected by GM composition. Notably, pronounced infections were readily achieved in inulin-fed mice given suppressive treatment for the full experimental period, suggesting that egg hatching and larval colonization was not impaired. However, we noted that whilst 16S copy number in faecal samples was reduced by at least several log-fold by antibiotic treatment in SSD-fed mice, indicating a successful suppression of the GM population, this reduction was more variable in inulin-fed mice (significant reduction in only one out of three studies) suggesting only partial depletion of the GM in these mice and a diet-antibiotic interaction (**Figure 5g**). Despite this, *Tm* burdens in inulin-fed mice were equivalent across experiments. Moreover, the antibiotic treatments drastically altered the GM composition in *Tm-*infected mice fed the different diets (**Figure 5h**; Supplementary Figure 3). In mice given the targeted antibiotic treatment, β-diversity analysis indicated that each treatment group formed a distinct cluster (adjusted *p* value<0.05 for all pairwise comparisons between treatment groups by PERMANOVA on Bray-Curtis distance metrics; **Figure 5h**). In the absence of antibiotics, the GM in SSD-fed, *Tm*-infected mice was dominated by the Bacteriodetes phylum, but in inulin-fed mice this changed to a dominance of Firmicutes, and a substantial expansion of Proteobacteria (mainly *Escherichia* spp. and *Parasutterella* spp.), consistent with our previous work^18^. However, in the presence of antibiotics, the GM was significantly restructured with the GM of the SSD-fed mice now dominated by *Streptococcus* spp. (Firmicutes) (**Figure 5h**, Supplementary Figure 3). Notably, the GM of the inulin-fed, antibiotic treated mice was considerably different to inulin-fed mice without antibiotics, with the dominant taxa now being *Enterococcus* spp. (Firmicutes), and *Bacteroides* spp. and *Parabacteroides* spp. (Bacteroidota) (**Figure 5h**, Supplementary Figure 3). Yet, worm burdens in inulin-fed mice were remarkably consistent despite these highly disparate GM compositions, indicating a dietary effect that was preserved despite significant perturbation of the underlying GM. We also profiled the GM composition of mice given continuous antibiotic treatment and where 16S copy number in faeces was significantly reduced (‘closed circles’ in **Figure 5g**). Here, GM composition significantly differed in mice fed either SSD or inulin without antibiotic treatment, but there were no differences between diets in antibiotic-treated mice (Supplementary Figure 4). Thus, the GM was effectively depleted by antibiotic treatment and normalized across the dietary conditions, yet *Tm* burdens remained high in inulin-fed mice, and negligible in SSD-fed mice (Supplementary Figure 4). Whilst the reasons for the discrepancies between the response to antibiotic treatment in different experiments remain elusive, our data thoroughly corroborate that the effect of inulin on *Tm* burdens is independent to host GM composition. Thus, these collective data demonstrate that GM-mediated metabolism of inulin plays a negligible role in *Tm* susceptibility. Instead, the observed effects may be attributed to the intrinsic properties of the dietary fibre.

### Removal of inulin from the diet during a critical window of immunity establishment rapidly restores *Trichuris muris* expulsion and attenuates inflammatory cytokine expression

Given that inulin-mediated effects on *Tm* burdens seemed independent of host GM composition, we postulated that the inulin fibre *per se* was mediating the effects on infection and host response. If this were the case, then removal of the inulin from the diet should rapidly restore resistance. Mice were fed either SSD or inulin until either 7 days or 14 days p.i. At this point, the diets were switched, until termination at day 21 p.i (**Figure 6a**). Strikingly, changing the diet at day 7 p.i. resulted in an almost complete reversal of the infection phenotype. Mice switched from SSD to inulin harboured substantial worm burdens at day 21 p.i, whilst those mice switched from inulin to SSD efficiently expelled their worms (**Figure 6b).** However, changing the diet at day 14 p.i. had the opposite effect; i.e., mice switched from SSD to inulin expelled their worms normally, whereas mice switched from inulin to SSD retained large numbers of worms (**Figure 6b**). These data suggest that there is a critical window between day 7-14 p.i. where inulin mediates its detrimental effects on host immune response and worm expulsion. This observation aligns well with data showing that in resistant mice, accelerated epithelial turnover responsible for *Tm* expulsion are established between day 7-14 p.i.^28^.

**Figure 6.**
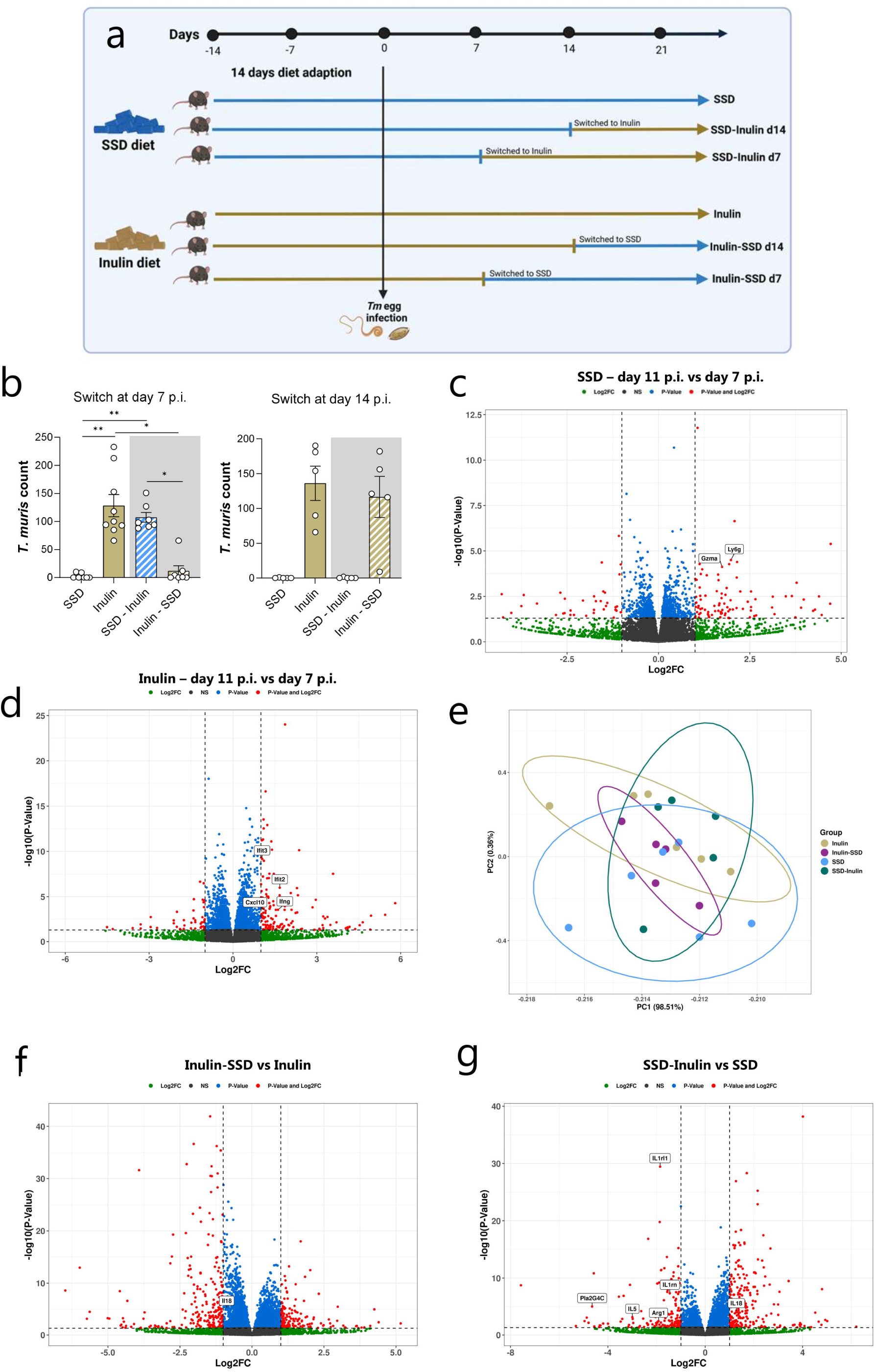
Effects of switching SSD and inulin diets on *Trichuris muris* burdens and caecal transcriptional responses. **a)** Schematic of experimental design. **b)** *Trichuris muris* (*Tm*) burdens at day 21 p.i. in mice fed SSD, inulin, or the diets switched at either day 7 p.i. or day 14 p.i. For day 7 switch, data are pooled from two independent experiments with (3-5 mice per group). For day 14 switch, data are from a single experiment with n=5 per group. **p>0.01; *p<0.05 by one-way ANOVA. **c-d)** Volcano plots showing differentially expressed genes (fold change >2; adjusted p value <0.05) identified by RNAseq at day 11 p.i. compared to day 7 p.i. in *Tm*-infected mice fed either SSD (**c**) or inulin (**d**). **e)** Principal complement analysis of the caecal transcriptome at day 11 p.i. of *Tm*-infected mice fed either SSD or inulin, SSD switched to inulin at day 7 .p.i., or inulin switched to SSD at day 7 p.i. **f-g)** Volcano plots showing differentially expressed genes (fold change >2; adjusted p value <0.05) identified by RNAseq at day 11 p.i. Mice were fed either Inulin, SSD, inulin switched to SSD, or SSD switched to inulin at day 7 .p.i.

To explore if the immune phenotype did indeed change within this ‘window of opportunity’, we profiled the transcriptomic response in whole caecum tissue during the diet-switch period. We focused on the period between day 7 p.i. and day 11 p.i. to establish a window of immune plasticity that could be altered by dietary intervention. First, we compared the caecal transcriptome at day 7 p.i. in mice fed either SSD or inulin, and their respective uninfected controls, which showed a distinct response in each dietary group, confirming our earlier data that the innate response to *Tm* is modulated by the inclusion of dietary inulin (Supplementary Figure 5). Notably, relative to infected mice fed SSD, infected, inulin-fed mice had increased expression of *Il18* and *Nos2*, confirming our results of exaggerated pro-inflammatory signalling early in infection. Moreover, infection in SSD fed mice enriched gene pathways related to IgE signalling, B-cell activity, and T-cell activation, relative to their-inulin fed counterparts (Supplementary Figure 5). Thus, already at this early time-point infected mice within the two dietary groups were established on different immune trajectories leading to either worm expulsion or retention.

Next, we investigated how the caecal transcriptome changed between day 7 p.i. and day 11 p.i. In SSD-fed mice, there were only modest changes between the two time points (12 genes differentially regulated with a fold change >2; adjusted p <0.05), including increased expression of *Ly6g* and *Gzma* at day 11 p.i. (**Figure 6c**). In contrast, there was a substantial change between the time points in inulin-fed mice with 85 genes being differentially regulated (fold change >2; adjusted p <0.05), including increased expression of *Ifng, Ifit2, Ifit3,* and *Cxcl10* at day 11 p.i., indicating that dietary inulin strongly favoured the development of a Th1 response within this time window (**Figure 6d**). To assess if removal of inulin from the diet attenuated this Th1 induction, we compared the transcriptome at day 11 p.i. in mice that were fed inulin for the whole period, and mice that were fed inulin until day 7 p.i. and then switched to SSD. Removal of dietary inulin for only those 4 days (day 7-11) was sufficient to resemble mice fed SSD for the whole period, suggesting that continual consumption of inulin fibre is imperative for Th1 dominated immunity, and that this rapidly (within 4 days) reverts to the ‘normal’ Th2 response upon cessation of dietary fibre stimulation (**Figure 6e**). Relative to mice fed inulin for the whole period, the expression of 278 genes (including *Il18*) was modulated (fold change >2; adjusted p <0.05) in mice switched to SSD at day 7 p.i. (**Figure 6f**). Conversely, switching SSD-fed mice to inulin at day 7 p.i. resulted in the differential expression of 227 genes (fold change >2; adjusted p <0.05) by day 11 p.i., including the upregulation of *Il18*. Downregulated genes included *Il1rn*, a negative regulator of the inflammasome, and numerous genes involved in the type-2 anti-helminth response, including *Il5*, *Arg1, Pla2g4c,* and the IL-33 receptor *Il1rl1* (**Figure 6g**). Collectively, these data strongly corroborate that the addition or removal of inulin from the diet results in rapid remodelling of the host response to infection in the caecum, with *Il18* expression as a key fulcrum.

## Discussion

The role of diet and the GM in resistance to enteric infection is increasingly recognized. Whilst there appear to be numerous benefits of dietary fibre, particularly in obese or diabetic individuals^40, 41^, reports of increased pathogen infection in mice fed diets rich in fermentable dietary fibre^42, 43^ begs the question of whether there may be important side-effects to high fibre intake. These would have important clinical implications, particularly in parts of the world where parasites or gut bacterial infections are widespread.

Here, we have shown that a key mechanism of inulin-mediated susceptibility to whipworm infection is dysregulated innate immune responses in the intestinal epithelium. Subsequently, altered immunometabolism (e.g. an increasing predominance of IDO1 activity) drives chronic infection. Suppression of innate pro-inflammatory cytokines such as IL-27 or IL-18 negated this rise in IDO1 and restored worm expulsion. It is interesting to note that, in the absence of infection, inulin-rich diets induced interferon signalling pathways, increased expression of antimicrobial peptides, and higher abundances of mucosal T-helper cells. How inulin causes these responses is not yet clear, but presumably the increased complexity of the fibre rich diet (relative to the very simple composition of SSD) was sufficient to induce a state of heightened immune reactivity in the gut. Indeed, substituting inulin for a synthetic AhR ligands was sufficient to raise *Tm* burdens. Whilst AhR ligation alone was not sufficient to recapitulate the effect of inulin, it does provide support for the notion that increased activation of host xenobiotic-metabolising pathways progressively down-regulates the highly skewed Th2 responses found in SSD-fed mice. Interestingly, a fibre-free diet (equivalent to SSD) in mice has been associated with heightened Th2 responses to food allergens^44^, which could align with our results of a stronger type-2 response to *Tm* infection in SSD-fed mice. Thus, highly simple diets, lacking any complex plant material, may predispose to type-2 inflammation, consistent with a role for dietary fibre in the prevention the development of allergies^45, 46, 47^.

Similar to inulin, infection with *Tm*, in the initial early stages, was also characterized by antimicrobial peptide production and interferon signalling. It could then be speculated that inulin should increase immunity to *Tm* by amplification of the same pathways upregulated by the host in response to the parasite. Instead, the combination of dietary inulin and *Tm* infection showed a strong antagonistic response, with downregulation of genes encoding mast call proteases, defensins, and goblet-cell derived effectors such as RELM-β. There is clearly an intricate diet-parasite interaction, as inulin *per se* does not drive a Th1 response, but only during *Tm* infection. Indeed, in other models, inulin fibre can worsen symptoms of type-2 inflammation associated with ulcerative colitis^48^, suggesting that suppression of Th2 immunity is not inherent to inulin, but specific for this model of whipworm infection. Thus, our data indicate an important context-dependent role of dietary fibres to regulate the innate response to an intestinal infection.

Inulin is a highly fermentable fibre, and the effects we describe here are not observed when non-fermentable fibres such as cellulose are incorporated into SSD^18^. A logical explanation would be that bacteria that thrive in inulin-rich environments, and/or the metabolites they produce, are responsible for creating a milieu that favours *Tm* persistence. However, our results tend to argue against this hypothesis, with high worm burdens apparent in antibiotic-treated mice, and FMT unable to confer susceptibility to infection.. We cannot rule out the effects of specific bacteria or associated metabolites, particularly as antibiotic treatment seemed to be less effective in inulin-fed mice, with *Bacteroides* spp. and *Parabacterioides* spp. (both of which are known to ferment inulin^49, 50^) still present in the GM following antibiotic exposure. However, GM-independent effects of dietary components on pathogen infection are not without precedent. Wolter et al. reported that chow diets favour the establishment of *Salmonella* or *Listeria*, relative to SSD, and that this is equivalent in both SPF and germ-free mice, suggesting that the physio-chemical composition of the diet *per se* is a key factor regulating host-pathogen interactions^51^. Indeed, whilst inulin is most-well known as a prebiotic fibre, increasing evidence suggests that inulin and other soluble fibres such as pectin (but not insoluble cellulose) are able to interact directly with host cells to regulate immune function, as evidenced by *in vitro* models and studies in gnotobiotic animals^14, 52, 53, 54^. Molecular docking studies also have reported that inulin could potentially bind TLR4^13^, providing a potential explanation for its Th1 dominated immunostimulatory effects. The rapid upregulation of *Il18* in response to inulin intake would be consistent with inulin directly activating a TLR-driven inflammasome process, and our successful restoration of immunity to *Tm* by blocking IL-18 *in vivo* provides a framework for future studies to disentangle the link between diet, innate immune function and infection.

In summary, our results describe a diet-immune-pathogen interaction whereby inulin fibre compromises immunity to intestinal whipworm infection. These data have clear translational implications, as dietary supplements which target the immune system and microbiome may need to be carefully evaluated in situations where enteric infections are commonplace.

## Methods

### Experimental animals, diet and infection

C57BL/6JOlaHsd female mice (Envigo, aged 6-7 weeks at start of experiments) were maintained in individually ventilated cages of 3-5 mice per cage. On arrival, mice were fed ad libitum either SSD (13 kJ% fat, ssniff Spezialdiäten, Germany), SSD with 10% long-chain inulin replacing corn starch, or SSD with added I3C (200 mg/kg), with free access to water (Supplementary Table 1). After two weeks diet adaptation, mice were either infected with 300 embryonated *Tm* eggs via oral gavage or served as uninfected controls. Technical details regarding parasite maintenance were as previously described^18^. Mice were sacrificed by cervical dislocation at day 3, 7, 11, or 21 post-infection (p.i.). All experimentation was conducted in line with the Danish Animal Experimentation Inspectorate (License numbers 2015-15-0201-00760 and 2021-15-0201-00831), and approved by the Experimental Animal Unit, University of Copenhagen according to FELASA guidelines and recommendations.

### Pharmacological interventions

For inhibition of IDO1 activity, mice received 2 mg/mL (pH 10.5) 1-methyl-DL-tryptophan (1-MT; Sigma Aldrich) in drinking water seven days prior to receiving *Tm* infection and until the end of the experiment (day 21 p.i.). 1-MT stock solutions were prepared in 100 mM NaOH and stored at -20°C. Drinking water preparations containing 1-MT or pH-adjusted (pH 10.5) control water were freshly prepared daily throughout the treatment period. For inhibition of AhR activity, mice were treated with the AhR antagonist CH-223191 (Sigma Aldrich) at a dose of 10mg/kg. Mice received daily oral gavage of CH-223191 in corn oil, or corn oil alone, from onset of *Tm* infection until the end of the experiment (day 21 p.i.). HD-5 was prepared as previously described^55^ and administered at a dose of 7.19 µg in PBS per mouse daily between day 0 and day 21 *Tm* infection. Control mice received PBS alone. For *in vitro* HD-5 experiments, *T. muris* larvae were recovered from inulin-fed mice at day 21. p.i., washed and then incubated in PBS at 37°C with or without HD-5 (10 µg/mL) overnight. The motility of the worms was then scored by an observer blinded to the treatment groups, as previously described^56^. For EdU treatment, mice were injected i.p. with EdU (10mg/Kg) 90 minutes prior to necropsy. For detection of EdU incorporation, cells were identified by fluoro-labelled azide using the Click-iT EdU Cell proliferation kit (ThermoFisher), according to the manufacturer’s instructions. Positive cells (5 colonic crypts/mouse) were visualized using a Zeiss AxioScope Fluorescent microscope, and images processed with ImageJ software (ImageJ, NIH, USA).

### Antibiotic Treatments

*Tm*-infected Mice were administered a cocktail of ampicillin (1g/L) and neomycin (1g/L) in drinking water from 4 days p.i. to 21 days p.i. Alternatively, where indicated mice received the ampicillin (1g/L), neomycin (1g/L) and colistin (35 mg/L) in drinking water from -7 days p.i. until day 21 p.i.

### In vivo cytokine depletion

Mice were treated with either 200 mg of purified anti-mouse IL6 (clone MP5-20F3; BioXCell), IL-27 (clone MM27.7B1; BioXCell) or IL-18 (clone YIGIF74-1G7; BioXCell), or respective isotype controls (clones RTK2071;BioLegend, eBM2a; Thermofisher; eBR2a; ThermoFisher) by i.p. injection every 4 days starting from day 0 (*Tm* infection) until day 21 p.i.

### Faecal GM transfers

Faeces was collected directly after defecation from mice that had consumed either SSD or inulin diets for 2-3 weeks. Faeces was immediately placed in airtight tubes containing PBS with 10% glycerol and stored at -80°C until needed. Donor faecal material was thawed and immediately gavaged into SSD-fed recipients. Where indicated, some recipient mice received an antibiotic cocktail (ampicillin and neomycin) for 7 days, and then were rested for 7 days prior to FMT. For real-time FMT studies, donor mice were fed SSD or inulin and infected with 300 *Tm* eggs. At defined time points during infection, faeces was collected from donor mice, placed in airtight tubes containing PBS, briefly vortexed, and then immediately gavaged into recipient mice that were fed SSD and at an equivalent time point of *Tm* infection.

### Metabolite analysis

Tryptophan and kynurenine levels in serum were assessed by LC-MS/MS. Ice-cold 100% methanol was added to thawed serum samples which were then centrifuged at 15,000 RCF at 5°C for 20 minutes. The supernatant was transferred to a clean tube, before evaporation using a SpeedVac Concentrator (SPD131DDA, ThermoFisher). Dried samples were reconstituted in 200 µL MilliQ water together with internal standards (Trp-d5 and Kyn-d4, final concentration 60 ng/mL). The samples were vortexed and filtered into HPLC glass vials, and then analzyed with a flow rate of 0.6 mL/min using a Zorbax 300SB-C8 column (150 mm length, 2.1 mm inner diameter and 5 µm particle size; Agilent Technologies, Denmark) connected toa Thermo Ultimate 3000 UHPLC coupled to Orbitrap Q Exactive MS (ThermoFisher) system. The mobile phase constituted0.1% v/v formic acid in MilliQ water (A) and 100% acetonitrile (B). The following gradient program was used to separate the analytes: 0.00 to 4.50 min: 5% B; 4.50 to 6.00 min: 5-90% B. The column was equilibrated for 4 min. The MS operating parameters were the same as described previously^57^.

### Histology and immunohistochemistry

Full thickness proximal colon tissue sections were taken at day 21 p.i. and fixed in 4% paraformaldehyde. Fixed tissue was paraffin embedded, sectioned and mounted onto glass slides, prior to staining. Sections were stained with either periodic-acid Schiff for goblet cell enumeration, or de-waxed and treated with citrate buffer for antigen retrieval prior to staining with rabbit anti-mouse polyclonal anti-indoleamine 2,3-dioxygenase antibody (Abcam, UK), and biotinylated anti-rabbit IgG (Abcam, UK).

### Cell isolation

Mesenteric lymph nodes were removed from mice and processed as previously described for flow cytometry^18^. Colons were dissected, stripped of fat and mesentery tissue, and flushed with cold PBS to remove mucus and intestinal contents. Colons were further cut into small segments and digested in HBSS supplemented with 10% FCS, 30 mM EDTA and 1 mM DTT (Sigma) on a shaking heat block for 20 min at 37°C and 900 rpm. IEC solutions were then filtered using 70 µM strainer and processed for RNA extraction. Lamina propria fractions were washed in PBS containing 5% FCS to remove EDTA solution and any remaining epithelial cells, then minced into smaller pieces using a scissors. The tissue was then digested twice with pre-warmed DMEM (Gibco) supplemented with Liberase (40 µg/mL; Roche) and DNase I (50 µg/mL) for 30 min each at 37°C and 650 rpm, with fresh digestion media added for each incubation. The resulting tissue was filtered through a 70 µM strainer, with cell pellets re-suspended and stored in DMEM with 5% FCS for subsequent flow cytometry.

### Flow cytometry

Mesenteric lymph node single cell suspensions were incubated for 20 min on ice with a combination of the following surface antibodies: anti-TCRβ (clone H57-597), anti-CD4 (RM4-5), anti-CD25 (PC61) anti-CD64 (X54-5/7.1), and anti-CD11b (M1/70) from BD Biosciences (Denmark); anti-I-A/I-E (M5/114.15.2), anti-CD103 (2E7) and anti-CD11c (N418) from Thermo Fisher Scientific (Denmark). For transcription factors, cells were fixed and permeabilized using the Foxp3 staining kit (BD Biosciences) following manufacturer’s instructions, and then incubated on ice for 30 min with the following: anti-Tbet (4B10) from BD Biosciences; anti-Foxp3 (FJK-16s), anti-Rorγt (B2D), and anti-GATA3 (TWAJ) from Thermo Fisher Scientific. Cells were analysed on a BD Accuri C6 cytometer (BD Biosciences), and data acquired using Accuri CFlow Plus software (Accuri Cytometers). Single cell suspensions isolated from lamina propria were stained with 4’,6-diamidino-2-phenylindole (DAPI) for dead cell exclusion following manufacturer’s instructions. Cells were then washed, and treated with anti-CD16/32 Fc block (BD Biosciences), before incubation on ice for 30 min with the following surface antibodies: anti-CD45 (30-F11), anti-TCRαβ (H57-597), anti-CD90.2 (30-H12) and anti-CD117 (2B8) from Biolegend (Nordic Biosite, Denmark); anti-CD4 (RM4-5), anti-CD8a (53-6.7) and anti-Nk1.1 (PK136) from BD Biosciences; anti-Nkp46 (29A1.4), anti-CD49b (HMa2), anti-CD11b (M1/70) and anti-FcεR1 (MAR-1) from Thermo Fisher Scientific. After fix/permeabilization using the FoxP3 kit cells were incubated for 20 min at room temperature with: anti-Tbet (O4-46) and anti-Rorγt (Q31-378) from BD Biosciences; and anti-GATA3 (TWAJ) from Thermo Fisher Scientific. Cells were analysed on a BD LSRII (BD Biosciences), and data acquired using FlowJo™ (BD Biosciences). Gating strategy is shown in Supplementary Figure 6.

### RNA extraction and quantitative PCR

Colonic epithelial cell fractions or whole caecum tissues were disrupted into a homogenous cell suspension using QIAzol lysis buffer using a gentleMACS Dissociator (Miltenyi Biotec). RNA was extracted using a RNAeasy mini kit (Qiagen) following manufacturer’s guidelines. Total RNA purity and concentration was assessed using a Nanodrop ND-100 spectrophotometer (Nanodrop Technologies). Quantitative PCR was conducted using PerfeCTa SYBR Green Fastmix (Quantabio) using the following program: 95°C for 2 min followed by 40 cycles of 15 s at 95°C and 20 s at 60°C. Primer sequences are listed in Supplementary Table 2. *Gapdh* was used as a reference gene for normalization and fold changes calculated using the ΔΔCT method.

### RNA sequencing

For isolated epithelial cells, RNA-seq (150bp paired-end) was carried out using the DNBSEQ-G400 platform (BGI, Copenhagen, Denmark). For caecal tissue, RNA-seq (150bp paired-end) was carried out using the Illiumina NovaseqX platform (BMKgene, Münster, Germany). Clean reads were aligned to the mouse genome and differential gene expression determined using DESeq2^58^ Pathway analysis was conducted using Gene Set Enrichment Analysis (Broad Institute, MA, USA).

### Amplicon sequencing for microbiota analysis

Prior to extraction, 750 μl ZymoBIOMICS™ Lysis Solution was added to faecal samples which were then lysed in a TissueLyser (Qiagen) at 30GHz for 10 minutes followed by centrifugation at 10,000 x g for 1 minute. Genomic DNA was extracted using the ZymoBIOMICS™ 96 MagBead DNA Kit (Zymo Research), following the manufacturer’s recommendations. Library preparation targeting the V3-V4 hypervariable region of the 16S rRNA gene and paired-end sequencing were conducted using the Illumina NovaSeq 6000 platform (Novogene, Cambridge, UK). Negative controls were sequenced with the samples and consisted of both extractions and PCR blanks.

### Bioinformatics and statistical analysis of microbiota

To assess the quality of the reads FastQC^59^ and MultiQC^60^ were used. The DADA2 pipeline^61^ was used to filter, trim, infer ASVs, merge forward and reverse reads, remove chimeric sequencing, and taxonomy assignment with the SILVA/v138 reference database^62^. Decontam was used to remove contaminant sequences^63^. The remaining ASVs were processed using the R-package Phyloseq^64^ and normalized using rarefaction. Microbiota composition was estimated using a PCoA with Bray-Curtis distance and statistical difference was estimated using PERMONOVA using the R-package Vegan^65^ and visualized using the R-package ggplot2^66^. All data analysis in R was performed using the R version 4.3.1.

### qPCR for 16S copy number

5 µL of extracted faecal DNA was added to 15 µL Master mix containing 1 µL forward primer (5’-3’ ACWCCTACGGGWGGCAGCAG), 1 µL reverse primer (5’-3’ ATTACCGCGGCTGCTGG), 3 µL water and 10 μl of RealQ Plus 2x Master Mix Green (Ampliqon 5000840-1250). The qPCR was run on a C1000 touch thermal cycler (BioRad 1851196) and CFX96™ Touch Real time PCR system (BioRad 105-5096), on the following program: A holding stage consisting of two minutes at 50°C, followed by 10 minutes at 95°C, then 40 cycles of 15 seconds at 95°C, 20 seconds at 58°C, and 40 seconds at 60°C. The 40 cycles were followed by the melting curve consisting of 20 seconds at 58°C, 40 seconds at 60°C, 15 seconds at 95°C, one minute at 60°C, 30 seconds at 95°C, and 15 seconds at 60°C. A standard curve was prepared using *E. coli* plasmid and copy number/ µL determined.

### Statistical analysis

P values of <0.05 were considered significant. Assumptions of normality were checked through Shapiro-Wilk tests, or inspection of histogram plots and Kolmogorov-Smirnov tests of ANOVA residuals. Parametric data were analyzed using unpaired t-tests, or one-way ANOVA or 2-way ANOVA followed by Tukey’s post-hoc testing, and presented as means ± S.E.M. Non-parametric data were analyzed using Mann-Whitney tests and results presented as median values. Details of each experiment are given in the appropriate figure legends.

## Acknowledgements

The authors are grateful to Cecilie Anastacia Stokkeby Koch, Mona Sharma, Mette Marie Schjelde, Emil Jakobsen, Dennis Sandris Nielsen, Denitsa Stefanova, Axel Kornerup Hansen, and Line Fisker Zachariassen for assistance. This work was supported by the Novo Nordisk Foundation (grants 0052422 and NNF22OC0074714) and Independent Research Fund Denmark (grant 2034-00245B).

## Author Contributions

Conceptualization - L.J.M., A.R.W.; supervision- M.N.L., S.M.T., M.T.L., B.A.H.J, A.R.W.; investigation, L.J.M., P.J., P.A., L.Z; experiment and data analysis – L.J.M., P.J., P.A., A.M.J., A.V.J., A.V.M., M.P., E.K., A.R.W; funding acquisition, A.R.W; writing & editing, L.J.M., B.A.H.J; A.R.W. All authors contributed to the final editing of the manuscript.

## Conflict of Interest Statement

The authors declare that the research was conducted in the absence of any commercial or financial relationships that could be construed as a potential conflict of interest.

## Data Availability Statement

Raw RNA-Seq data are available at GEO under accession numbers GSE284172 and GSE281400. Raw 16S amplicon sequencing data are available at ENA under accession number PRJEB89498. All other data are contained with the manuscript or the supplementary files.

## Notes

https://www.ncbi.nlm.nih.gov/geo/query/acc.cgi?acc=GSE284172

https://www.ncbi.nlm.nih.gov/geo/query/acc.cgi?acc=GSE281400

https://www.ebi.ac.uk/ena/browser/view/PRJEB89498

### Competing Interest Statement

The authors have declared no competing interest.

