## Supplementary Material for "Dietary fibre promotes chronic whipworm infection through direct and time-dependent modulation of innate immunity"

**Contents:**

- Supplementary Tables 1-2
- Supplementary Figures 1-6

**Supplementary Table S1:** Diet composition of semi-synthetic diet and inulin-supplemented diet. All diets were purified and pelleted. Purified long-chain inulin contained average degree of polymerization (DP) ≥ 23.

| Feed component (g/100g) | SSD | Inulin |
|---|---|---|
| Casein | 20.0 | 20.0 |
| Corn starch | 37.6 | 27.6 |
| Inulin[1] | - | 10.0 |
| Maltodextrin | 15.0 | 15.0 |
| Sucrose | 10.0 | 10.0 |
| Cellulose powder | 5.0 | 5.0 |
| L-Cysteine | 0.2 | 0.2 |
| Vitamin premix | 1.0 | 1.0 |
| Mineral & trace element mix | 6.0 | 6.0 |
| Choline chloride | 0.2 | 0.2 |
| Soybean oil | 5.0 | 5.0 |
| **Crude Nutrients (%)** | | |
| Crude protein | 17.9 | 17.6 |
| Crude fat | 5.1 | 5.1 |
| Crude fiber | 5.0 | 14.4 |
| Crude ash | 5.4 | 5.4 |
| Starch | 36.2 | 26.6 |
| Sugar | 11.0 | 11.1 |
| **Energy (MJ [or kcal] ME/kg)** | 15.4 | 13.9 |

[1] Long-chain inulin purified from chicory root (Orafti HP, Beneo, Netherlands).

**Supplementary Table S2:** Primers used for qPCR

| Gene | Forward Primer (‘5-‘3) | Reverse Primer (‘5-‘3) |
|---|---|---|
| *Ahr* | AGTTCTTGTTACAGGCGCTGA | GCCCGGTCTTCTGTATGGAT |
| *Ang4* | TAGACTCGTCCCCAGTTGGA | CTGAGCCAGAGTTGGAGGAAT |
| *Defa5* | TCAAAAAAGCTGATATGCTATTG | AGCTGCAGCAGAATACGAAAG |
| *Gapdh* | TATGTCGTGGAGTCTACTGGT | GAGTTGTCATATTTCTCGTGG |
| *Ifng* | GACTGTGATTGCGGGGTTGTA | TCACTGCAGCTCTGAATGTTTCT |
| *Il13* | GGCAGCATGGTATGGAGTGT | CTTGCGGTTACAGAGGCCAT |
| *Il33* | CGGCAGAATCATCGAGAAACCT | GCCGGGGAAATCTTGGAGTTG |
| *Mcpt2* | GGCAAAATGCAGGCCCTACT | TCCTTCGAACCGTTCTTAGTGG |
| *Retnlb* | CTGATAGTCCCAGGGAACGC | GTCTGCCAGAAGACGTGACA |
| *Tslp* | AGGGGCTAAGTTCGAGCAAA | TTTTGTCGGGGAGTGAAGGG |

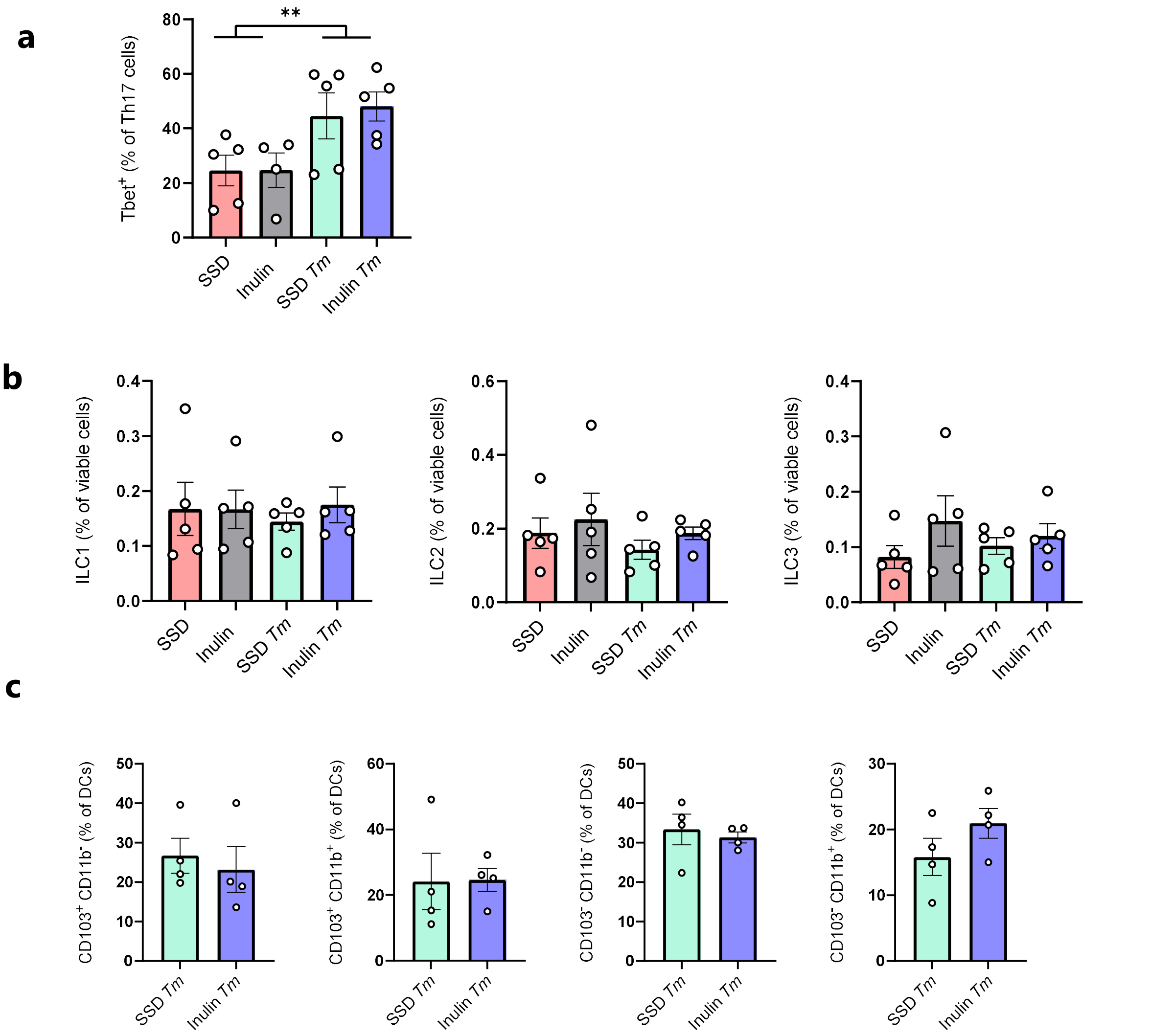


### Supplementary Figure 1

a) Proportions of Tbet+ cells withing Th17 population in large intestinal tissue, b) proportions of of type-1, type-2 and type-3 innate lymphoid cells (ILC) in large intestinal tissue, and c) proportions of dendritic cell (DC) subsets in mesenteric lymph nodes, in mice fed either SSD or inulin, with or without 7 days of *Trichuris muris* (*Tm*) infection. DCs were defined as $CD11c^+$, $CD64^-$, $MHCII^+$. n=5 per treatment group, shown is mean ± S.E.M.

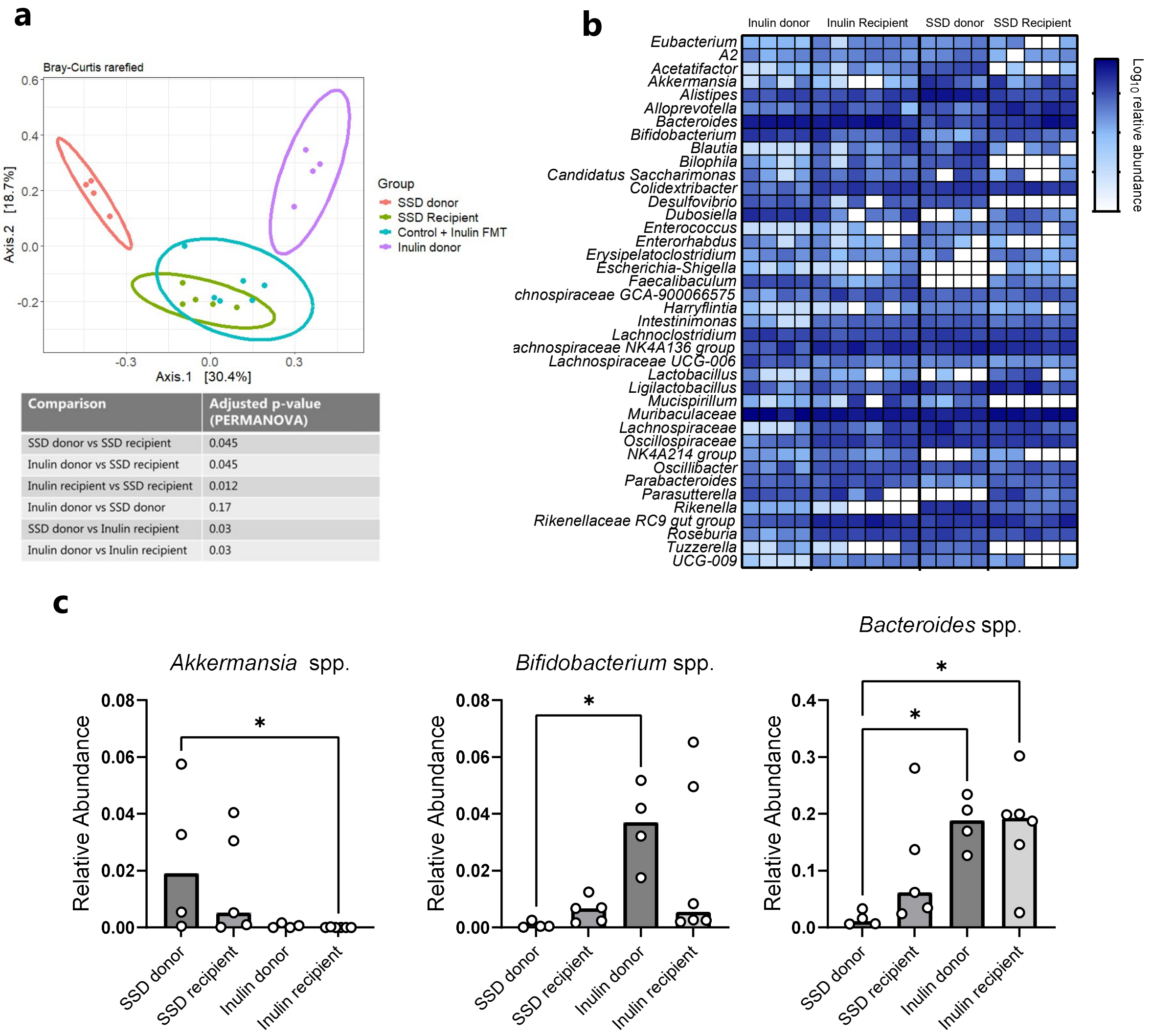


| Comparison | Adjusted p-value (PERMANOVA) |
|---|---|
| SSD donor vs SSD recipient | 0.045 |
| Inulin donor vs SSD recipient | 0.045 |
| Inulin recipient vs SSD recipient | 0.012 |
| Inulin donor vs SSD donor | 0.17 |
| SSD donor vs Inulin recipient | 0.03 |
| Inulin donor vs Inulin recipient | 0.03 |

### Supplementary Figure 2

a) PCoA analysis (Bray-Curtis distance metrics) of faecal microbiota of SSD- or inulin-fed donor mice, and recipient mice prior to infection. n= 4-6 per treatment group. Shown also are PERMANOVA comparisons between groups.

b) Relative abundances of genera (and *Muribaculaceae, Lachnospiraceae,* and *Oscillospiraceae* familes) in donor and receipeint mice.

c) Relative abundances of selected genera. *$p$ <0.05 by Kruskal-Wallis test and Dunn's post-hoc testing.

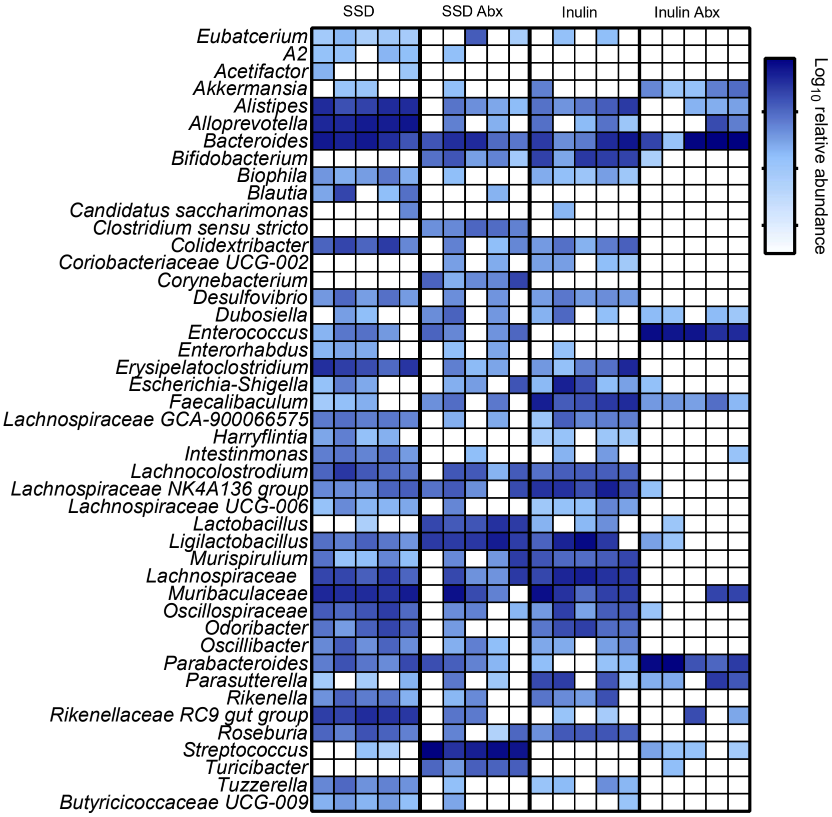


### Supplementary Figure 3

a) Relative abundances of genera (and *Muribaculaceae, Lachnospiraceae,* and *Oscillospiraceae* familes) in *T.muris*-infected mice treated with either a targeted antibiotic dose (day 4 p.i. to day 21 p.i.), or no antibiotic treatment, and fed either SSD or inulin.

**a**

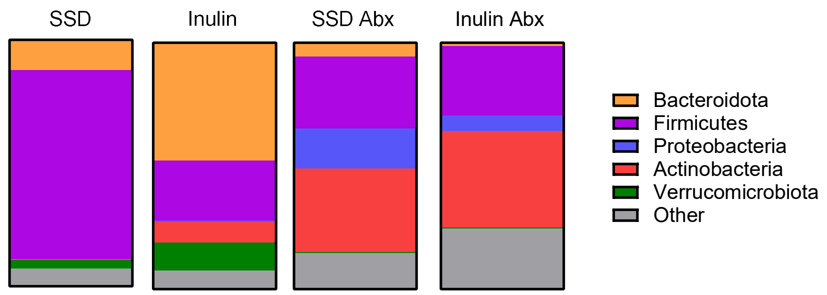


**b**

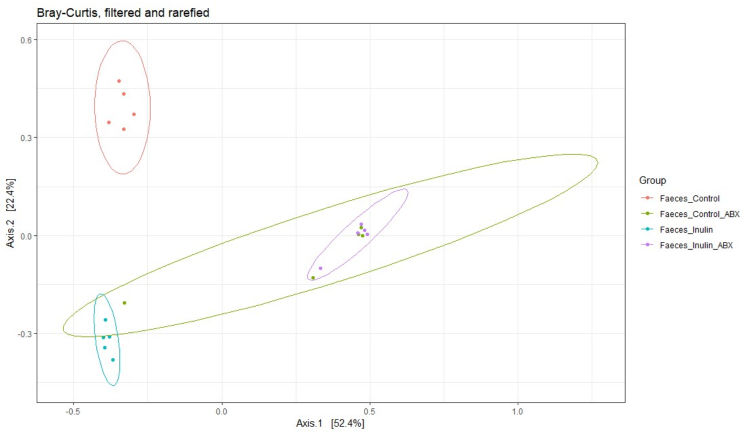


| Comparison | Adjusted p-value (PERMANOVA) |
|---|---|
| SSD vs Inulin | 0.046 |
| SSD vs SSD+Abx | 0.045 |
| SSD vs Inulin+Abx | 0.045 |
| Inulin vs SSD+Abx | 0.046 |
| Inulin vs Inulin+Abx | 0.049 |
| SSD+Abx vs Inulin+Abx | 1 |

### Supplementary Figure 4

a) Relative abundances distributed by phylum in mice infected with *Trichuris muris* for 21 days and fed either control semi-synthetic diet (SSD), inulin, with or without antibiotic (Abx) treatment for the full experimental period.

b) PCoA analysis (Bray-Curtis distance metrics) of faecal microbiota of control (SSD)- or inulin-fed mice, with or without antibiotic (Abx) treatment for the full experimental period. n= 5 per treatment group. Shown also are PERMANOVA comparisons between groups.

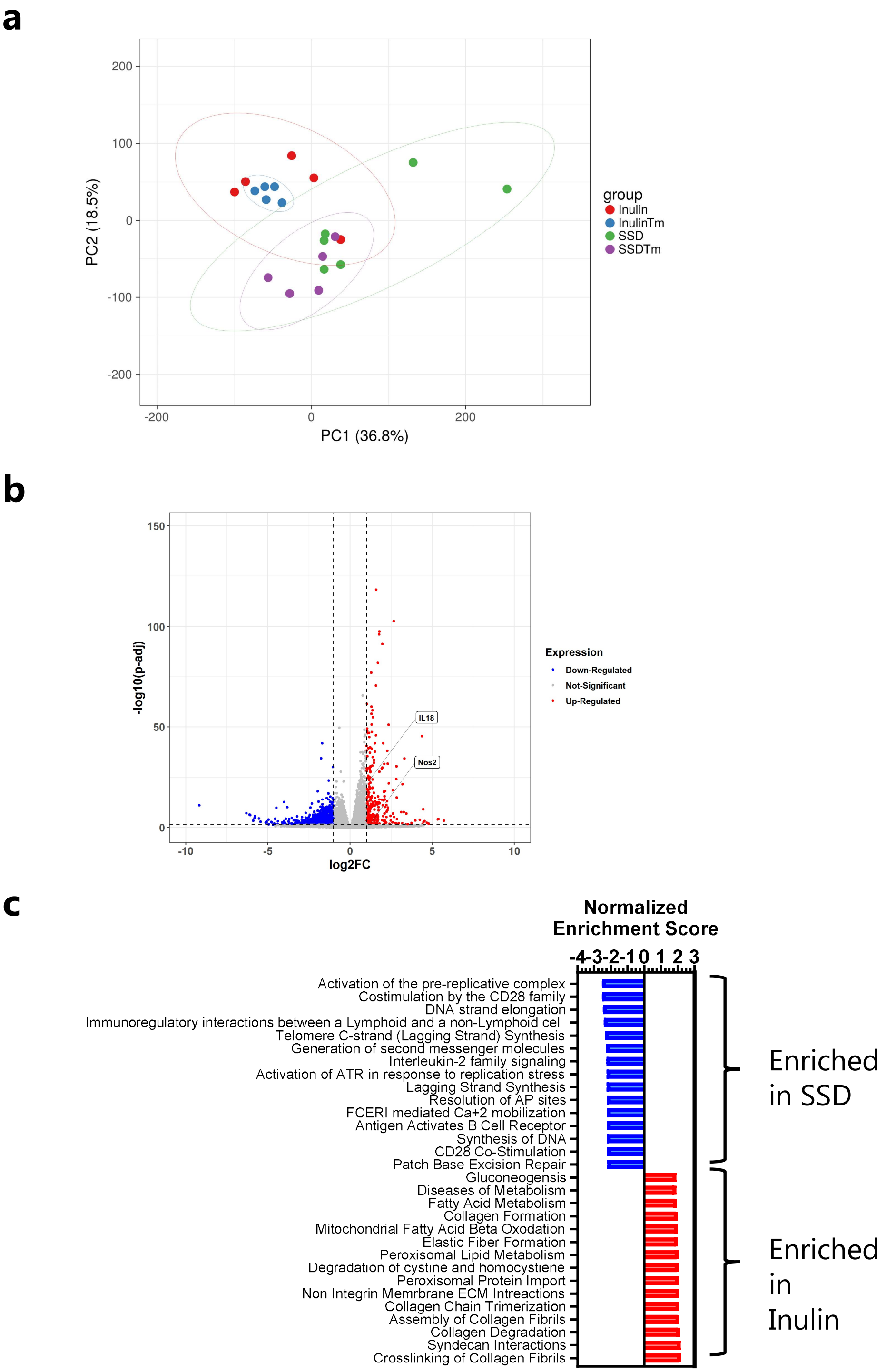


### Supplementary Figure 5

a) Principal component analysis of the caecal transcriptome in mice fed either SSD or inulin, with or without 7 days *Trichuris muris* (*Tm*) infection. N= 5-6 per group

b) Significantly regulated (fold change >2; adjusted *p* value <0.05) in *Tm*-infected mice (day 7 p.i.), fed with inulin, relative to infected mice fed SSD.

c) Significantly regulated gene pathways (q<0.05) identified by gene set enrichment analysis in *Tm*-infected mice (day 7 p.i.), fed either SSD or inulin

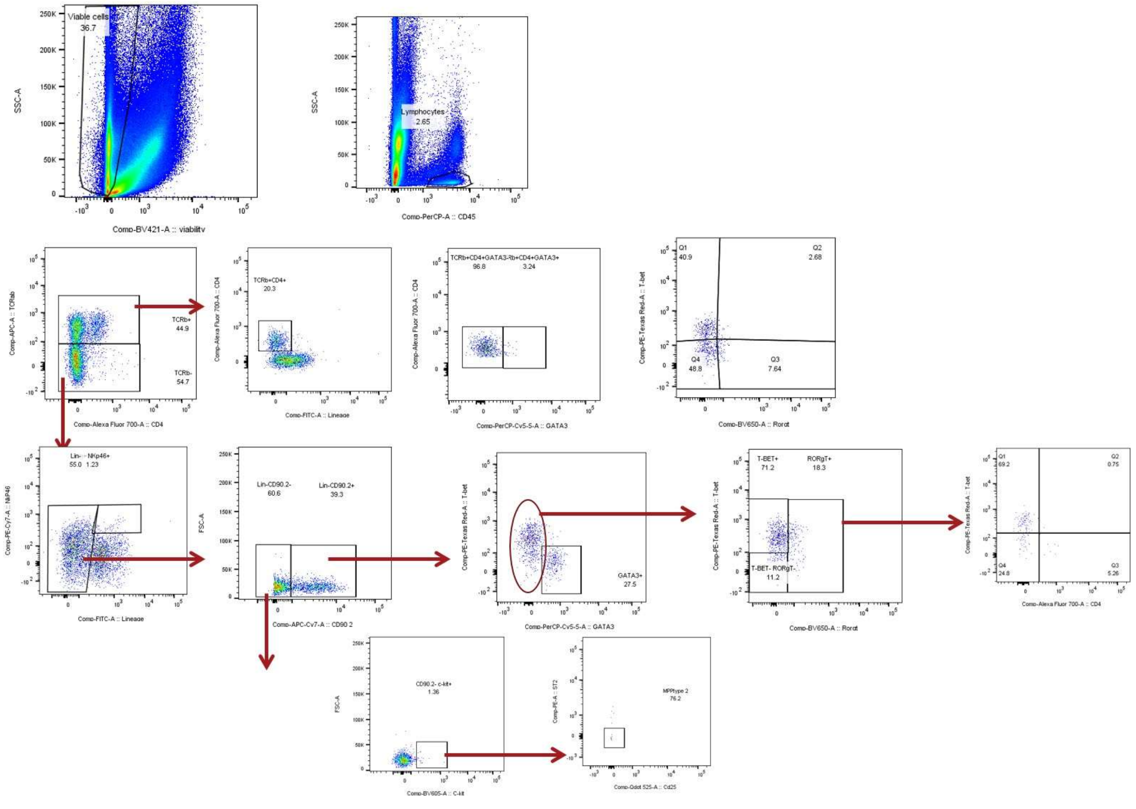


### Supplementary Figure 6

Gating Strategy for identification of T-cells and innate lymphoid cells in intestinal tissue.
